# Adoption and consequences of new light-fishing technology (LEDs) on Lake Tanganyika, East Africa

**DOI:** 10.1101/619007

**Authors:** Huruma Mgana, Benjamin M. Kraemer, Catherine M. O’Reilly, Peter A. Staehr, Ismael A. Kimirei, Colin Apse, Craig Leisher, Magnus Ngoile, Peter B. McIntyre

## Abstract

Maintaining sustainable fisheries requires understanding the influence of technological advances on catch efficiency. Fisheries using light sources for attraction could be widely impacted by the shift to light emitting diode (LED) light systems. We studied the transition from kerosene lanterns to LED lamps in Lake Tanganyika, East Africa, examining factors that led to adoption as well as the impact of the new light sources on fish catch and composition. We used a combination of field experiments with catch assessments, fisher surveys, underwater light spectra measurements, and cost assessments to evaluate the impact of switching from kerosene to LED lamps. Overall, we found a very rapid rate of adoption of homemade outdoor LED light systems in Lake Tanganyika. Most of the batteries used to power these lamps were charged from the city power grid, rather than photovoltaic cells. The LED light spectra was distinct from the kerosene light and penetrated much deeper into the water column. Regardless of light type, most of the fish caught within the two dominant species were below maturity, indicating that current fishery is not sustainable. Although the LED lamps were associated with a slight increase in catch, environmental factors, particularly distance offshore, were generally more important in determining fish catch size and composition. The main advantages of the LED lamps were the lower operating costs and their robustness in bad weather. Overall, the use of battery-powered LED lighting systems to attract fish in Lake Tanganyika appears to reduce economic costs but not contribute new impacts on the fishery.

## Introduction

Fish are a critical natural resource, and artisanal fisheries provide a key source of protein to many people around the world (1). If managed properly, fisheries can be considered a renewable resource (2), but advances in technology periodically allow for new access and extraction, and require reassessment of sustainable harvesting. Many changes related to the adoption of new technologies in artisanal fisheries are not documented nor properly assessed, and thus cannot be properly integrated into management strategies (3).

An important natural resource, the pelagic fishery in Lake Tanganyika, East Africa, faces a number of challenges. This critical fishery substantially contributes to poverty reduction (annual earnings of USD 10 million or greater) and provides food security in all riparian countries (Tanzania, Burundi, The Democratic Republic of Congo, and Zambia). Explicitly, the lake’s pelagic fish catches are vital source of food and livelihoods to millions of people living in the lake basin (4, 5). Presently, fishing pressure is very high with declines in catch per unit effort in some areas of the lake and the potential for local overfishing (5-7). There are also some indications of destructive fishing methods due to increased presence of illegal fishing gears (8). These issues of fisheries conservation and sustainability are widespread, as similar challenges exist in the other East African Great lakes as well as other inland lakes and reservoirs(9, 10).

Understanding the impacts of changes in fish gear is broadly important for many African lakes, and light attraction is one of the essential factors in pelagic fisheries (3, 11-13). Historically, fishermen used fire to create light to attract and aggregate fish at night for catching them by lift net, eventually switching to kerosene presume lamps (4, 14, 15). Recently, however, there has been a switch in the lighting gears from kerosene to light emitting diode (LED) lamp systems, and the effects of this on catch size and composition is unknown.

This study aimed to assess the costs of potential consequences of switching from using kerosene lanterns to various types of LEDs systems in Lake Tanganyika. Our overall aim was 1) to understand factors that influence the adoption of this new LED lighting technology and 2) to determine whether LED systems influence overall fish catch and composition.

## Material and methods

This study was conducted on Lake Tanganyika, in the northwestern region of the lake near Kigoma, Tanzania (Fig 1). Fishing is an important part of the community; in the Kigoma area in 2011, there were approximately 10,600 fishermen and 4,800 fishing crafts respectively of which 27% were lift net fishing units employing approximately 7,800 fishermen as crew (National Coordination Unit, 2011). The lake’s pelagic fishery is composed of six species, two endemic clupeids *Stolothrissa tanganicae* and *Limnothrissa miodon*, and four endemic centropomids of the genus *Lates* (4, 16) (*Lates angustifrons*, *Lates mariae*, *Lates stappersii*, and *Lates microlepis*). Of these, the clupeids (’dagaa’) and *L. stappersii* (’migebuka’) are the dominant part of the catch.

**Fig 1.**
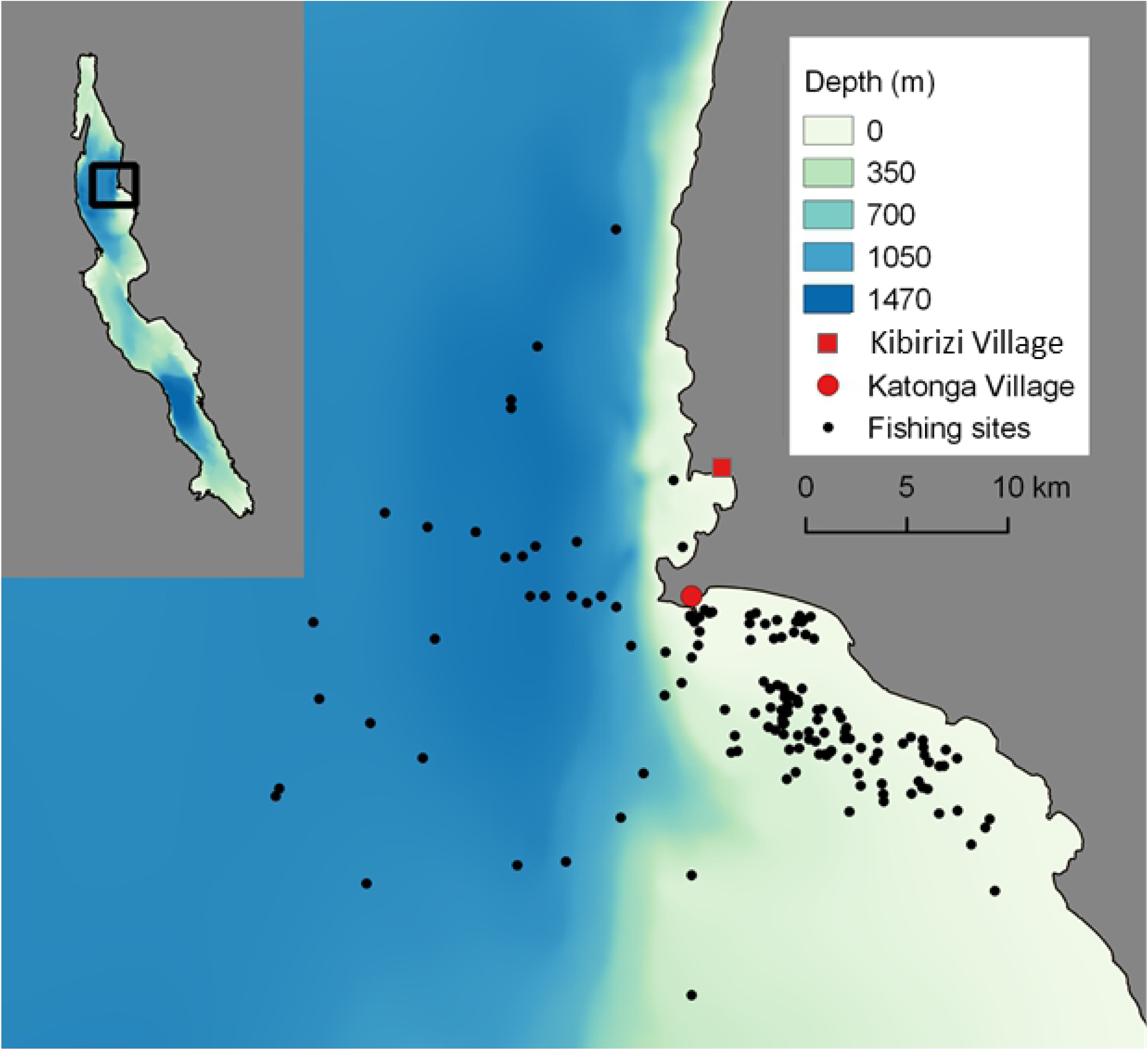
Map of the study site near Kigoma Bay, Tanzania on Lake Tanganyika. The adoption survey of fishermen was conducted at Kibirizi and Katonga villages. The assessment of fish catches was done at Katonga village. Colours represent water depth in the region. Circle size represents size of the catch. Inset shows the region of the study in northwestern Lake Tanganyika.

Artisanal fishing in the pelagic zone is conducted using liftnets (6-8 mm mesh) with artificial light during dark nights (3, 4, 15). Catch is dominated by clupeids (70-80%) followed by *L. stappersii* (5-15%), with by-catches of cichlids, catfishes and large *Lates* (3). Inshore areas of the lake are known to be breeding grounds for clupeids and nursery for some *Lates* species (3, 4), as well as permanently habited by cichlid fishes (17), so catch composition is influenced by fishing location.

In order to establish influence of lamp types on catches, three similar fishing units at Katonga were used to compare catches among traditional kerosene lamps (’karabai’) and two types of LED lamps, with lamps rotating monthly among units from the end of the dry season (5 October 2015) into the early wet season (1 January 2016). The first LED lamp commonly being used are outdoor LED lights (Model: B-10W) purchased locally and combined into a multi-light system called ‘spesho’ (hereinafter, LED-S). The second LED system was a commercially available LED system manufactured by Rex Energy (hereinafter, LED-R) (bulbs; Hella Sea Hawk XL 12/2W LED 0950 series), which is currently becoming available in local markets. The wattage per meter squared was similar among lamp types and fishing units always set more than a kilometre apart.

We measured the total weight of the daily catch using a field hanging scale. One to three homogenized handfuls of the fish sample (approximately 0.5-1.0 kg) were used for species composition analysis. Individual fishes from each taxon were weighed to the nearest 0.1 gram using a top-loading digital scale (Huazhi Sai Xijie electronic balance, model; TP-A1000), and their total lengths (TL) measured to the nearest millimetre from the tip of the mouth to the largest caudal ray.

To assess whether there were differences in how deep the light from the different lamps penetrated into the water column, we measured the spectral properties of each lamp at night. A submersible Stellarnet BLACK-Comet-SR Spectrometer (model, BLK CXR) measured light from 400 to 700 nm at depth (5-467 cm) and we used standard exponential light attenuation models to calculate the depth, which a single photon of light would reach. A questionnaire survey was conducted to assess the rapidity of the switch from kerosene lamps to LED lamps at two landing sites near Kigoma, Kibirizi to the north and Katonga to the south.

We collected some of the main environmental parameters known to influence fish catch such as moon phases, fishing distance from the shore (that correlates with water depth) and wind speed (3). The distance of each fishing site from shore was determined from GPS locations provided by fishers and calculated using QGIS. On nights when GPS data were not available, we estimated fishing location from travel time and bearing, as recorded by the boat captain using a Suunto manual diving compass (model SK4). Average wind speed was calculated for the period of fishing using measurements from an onshore weather station located at TAFIRI-Kigoma. The weather instrument used was Vantage Pro2 (model: 6162EU, Davis Instruments, USA) located at a height of 12 m above the ground. The potential moonlight was estimated as the average moon fullness fraction integrated over the hours when fishing lamps were used, based on moon fullness calendars (http://aa.usno.navy.mil/data/docs/MoonFraction.php), and moonrise/moonset times (http://aa.usno.navy.mil/data/docs/RS_OneDay.php). We refer to this as “potential” moonlight because it does not account for cloud cover.

A Boosted Regression Tree (BRT) statistical model was used to factor out the potential influence of environmental variables. The BRT model is a combination of regression decision trees and a boosting algorithm that can account for non-linear relationships between predictor variables and the response, missing predictor data, and interactions between categorical and numeric predictor variables (18). BRT analysis was completed using the “dismo”(19) and “gbm”(20) packages in R. All BRTs were fit with a model complexity value of “5” allowing for high levels of interactive effects between predictor variables. After fitting the BRT model, we used it to remove the influence of all variables other than lamp type on total daily catches. We did this using the equation

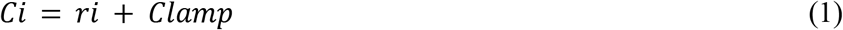

Where *C*_*i*_ is the i^th^ observation of total catches with variability attributed to all other variables other than lamp type removed, *r*_*i*_ is the model residual associated with the i^th^ observation, and *C*_*lamp*_ is the modelled estimate of total daily catches estimated for each lamp type holding all other predictor variables at their median. We used a similar approach to examine influence on fish size for the three main species.

## Results

There was rapid adoption of LED systems for night fishing. All respondents (n=26) who were interviewed were from Kibirizi and Katonga landing sites. The composition constituted boat crews (n=17), boat owners (n=8), and boat owners who practiced fishing by themselves with other crewmembers (n=1). All respondents acknowledged having used LEDs and kerosene lamps at some point in their lifetime as light sources for fishing. The shifts towards using LED lamp systems began in 2010 (Fig 2). Between 2010 and 2012, fishermen used an improvised LED system locally called ‘umua’, as the outdoor LED bulbs now used to build LED lamps (’spesho’) were not yet available. ‘Umua’ are made by dismantling circuits from LED flashlights and then attaching these inside hemispheres of big stainless steel bowls or housings of outdoor flood lights. The major shift happened after 2014 when outdoor LED bulbs became available, at which point more than 90% (n=24) of the respondents were using LED lamps. All respondents indicated that they no longer use the pressurized kerosene and have switched to homemade ‘spesho’ LED lamps between 10 to 22 LED bulbs rated between 3 - 10 watts; nobody used manufactured LED lamps (LED-R). Most respondents (n=17) believed that LED provided them with better catches.

**Fig 2.**
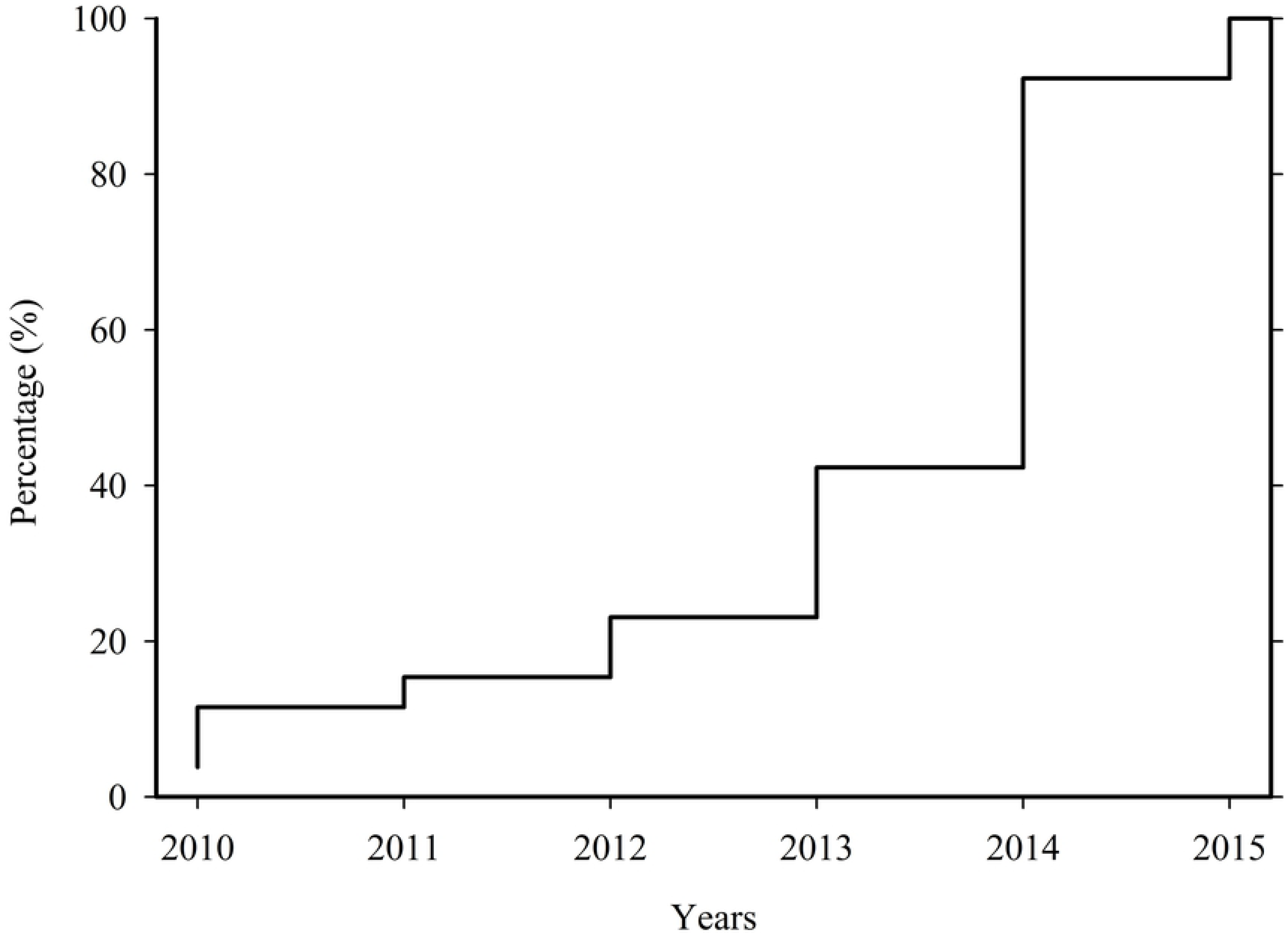
Cumulative percentage (%) of fishers switching from kerosene lanterns to LED-based lighting systems, showing the rapid adoption of this technology.

The operating costs were cheaper for LED systems relative to kerosene lamps. A single kerosene lamp requires 1-2 litres of kerosene daily (US $ 0.92 per litre during this period) as well as other replacement parts (US $ 9.0 per month). As fishing is typically done 20 days per lunar cycle, the monthly operating cost for one kerosene lamp is approximately US $50. In contrast, LED systems had monthly costs of $4-6 with daily battery charging. Batteries were charged at established kiosks at the landing sites and cost around US $0.92-1.40 per day. Typically, the charge lasts one to two nights of fishing, and each fishing unit uses two batteries (i.e. one battery per fishing boat). Most of the batteries being used to run the LED systems were being charged from the city main power supply (73%) (Fig 3). Only 15% of the respondents used solar panels, while 12% alternated between city power and solar panel charging kiosks to charge their batteries during the rainy season.

**Fig 3.**
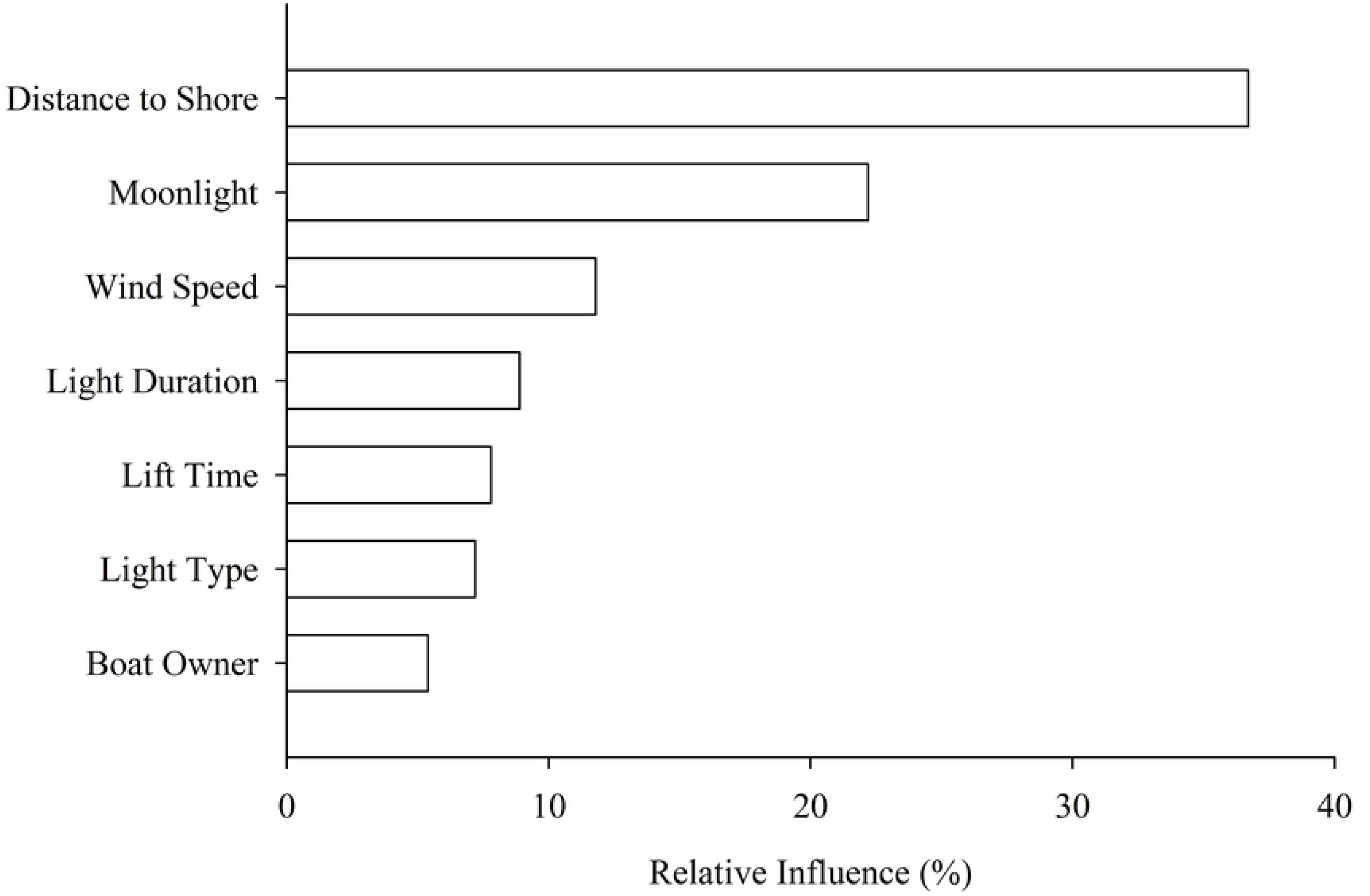
The relative importance of 7 drivers of fish catch variability. While the kind of light used has a small effect on fish catches, this effect is outweighed by other drivers of fish catch variability.

The adoption of LED lamps has been straightforward. Fishers did not require changes the size and dimensions of most of their accessories used in fishing, including the lift-net sizes and the lengths of the poles for hauling the net and holding the lamps. Boat size and number of fishermen have remained the same as when kerosene lamps were being used, and the depth to which they deploy their liftnets was still the same (around 100 m).

In terms of light energy emission, the LED lamps outperformed kerosene lamps. Light attenuation was greatest for kerosene lamps (K_d_ = 1.66 m^−1^), followed by the manufactured (LED-R) and homemade ‘spesho’ (LED-S) systems. (K_d_ = 0.53 m^−1^ and 0.46 m^−1^ respectively). Correspondingly, the depth at which one photon of light would be present from each lamp type was deepest for LED-S (125 m), followed by LED-R (120 m), and considerably shallower for the kerosene light (70 m).

Catches with LED lamps were slightly higher than the catches with kerosene lamps, but variation was explained primarily by environmental conditions (Table 1). Total daily catches ranged from 0 to 1476 kg of fish per boat per lamp type per night with a mean catch of 76 kg and a median catch of 21 kg. For total catch, the BRT model performed well as observed and modeled data were highly correlated (r = 0.79, Kendall rank correlation) with a median absolute error of 1.95 kg. The model also revealed that lamp type had a weak average effect on the total daily catches of fish per boat pair compared to the other predictors in the BRT model (relative influence = 8.2%) (Table 1). The distance from the shoreline, moonlight, and wind speed were found to be influential factors in the amount of fish caught per day per fishing unit, where their relative influences were 36.7, 22.2 and 11.8% respectively (Fig 3). Distance from shore was important up until about 5 km, beyond which it did not strongly influence catch. The duration of lighting, time net was hauled, and lamp type were less important when compared with the previous three environmental variables (relative influence of 8.9, 7.8 and 7.2% respectively; Fig 3). Within the lamp types, fishers using ‘spesho’ LED-S (30.9%) and manufactured LED-R (66.0%) lamps caught more fish as compared to fishers who were using kerosene lamps. Finally, boat owners had no influence on the daily catches.

**Table 1:**
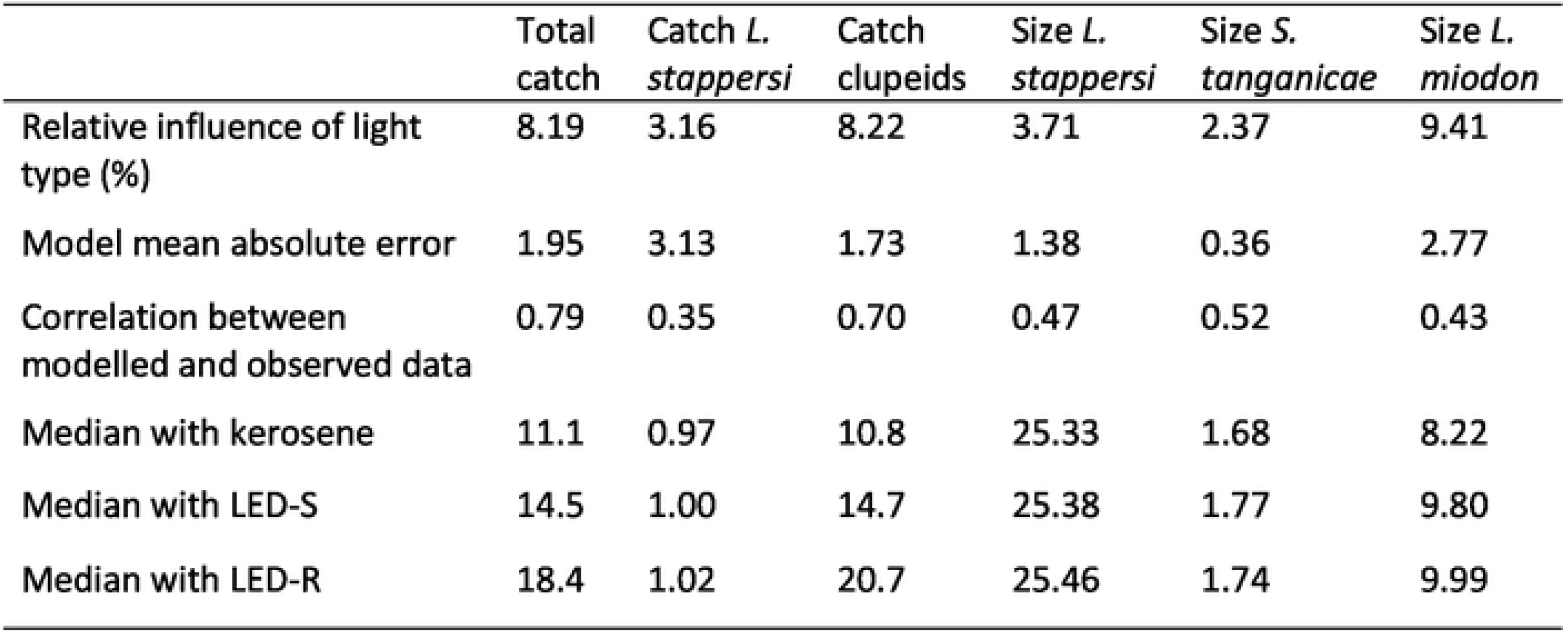
Results from the Boosted Regression Tree analyses showing differences among the lamp types for fish catch and size. Units are kg for catch, g for fish size.

|  | Total<br>catch | Catch *L.*<br>*stappersi* | Catch<br>clupeids | Size *L.*<br>*stappersi* | Size *S.*<br>*tanganicae* | Size *L.*<br>*miodon* |
| --- | --- | --- | --- | --- | --- | --- |
| Relative influence of light<br>type (%) | 8.19 | 3.16 | 8.22 | 3.71 | 2.37 | 9.41 |
| Model mean absolute error | 1.95 | 3.13 | 1.73 | 1.38 | 0.36 | 2.77 |
| Correlation between<br>modelled and observed data | 0.79 | 0.35 | 0.70 | 0.47 | 0.52 | 0.43 |
| Median with kerosene | 11.1 | 0.97 | 10.8 | 25.33 | 1.68 | 8.22 |
| Median with LED-S | 14.5 | 1.00 | 14.7 | 25.38 | 1.77 | 9.80 |
| Median with LED-R | 18.4 | 1.02 | 20.7 | 25.46 | 1.74 | 9.99 |

After removing the variability in *L. stappersi* catches attributed to other factors, fishers using LED-S and LED-R lamps caught more *L. stappersi* on average (about 2.9% and 5.2% more, respectively) than fishers using kerosene lamps (Table 1). The BRT model for *L. stappersi* total catches performed poorly with observed and modelled data weakly correlated (r = 0.35, Kendall rank correlation) and a median absolute error of 3.13 kg. Lamp type had a weaker than average effect on the total daily catches of fish per boat pair compared to the other seven predictors in the BRT model (relative influence = 3.2). Lamp type accounted for only 4.1-7.6 % of the variation in average fish size explained by boosted regression trees.

Overall, the clupeids *S. tanganicae* and *L. miodon* made up the bulk of the fish biomass caught during the experiment by contributing about 87% of the total catches (Fig 4). *L. stappersii* were also common but rarely dominated the total catches (12%). The remainder of the catch was composed of a range of species. These included the killifish *Lamprichthys tanganicae* and the deep-water cichlid *Bathybates* that feeds on pelagic species. *Chelaethiops boulenger* is a pelagic cyprinid found in the Malagarasi River and the pelagic and coastal zones of Lake Tanganyika. Moreover, although rarely, nearshore demersal fish species (mainly cichlids) were also caught. Occasionally, pelagic shrimp ranging from 1-2 cm long were found in the fish catches and some sponges that are commonly found on soft-bottom in the lake were also recorded.

**Fig 4.**
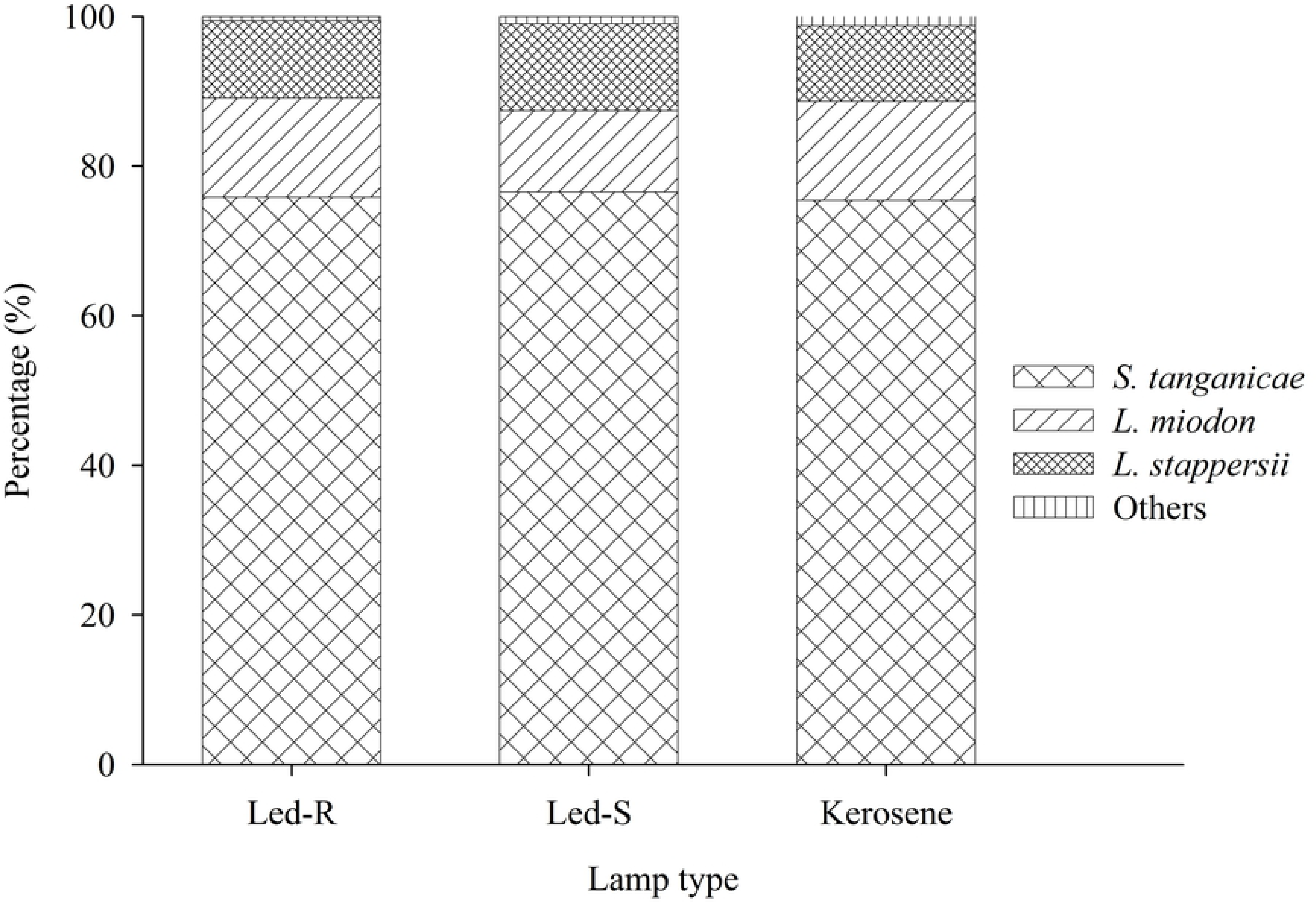
Species composition by weight of all fish caught in the study as a function of fishing lamp type.

## Discussion

The rapid adoption of LED lamps was driven by the relatively low operating costs compared to kerosene lamps. Our estimate of about US$50 per month operating costs for the use of a single kerosene lamp is similar to the cost found for the East African region by (21). The operating costs of LED lamps were eight to twelve times lower than kerosene lamps. For a complete economic analysis, the lifetimes and costs associated with the batteries would also need to be incorporated. The adoption of the LED lamps has led to the replacement of kerosene-selling kiosks with businesses associated with battery charging powered by city grid or diesel generators in remote areas. There are sustainable ways of charging LED lamp batteries by using solar panels (21, 22), but the initial investment cost in solar panels is high for an individual fishing team. The shift toward outdoor LEDs bulbs began in 2013, when they became available in local shops around Kigoma at a reasonable price (US$5 per bulb). The ability to design lower-cost, locally appropriate LED lamps is probably one of the contributing factors for the adoption (21), as LED systems (such as LED-R) made for specifically for fishing are expensive (US$175) relative to what is earned from fishing (21).

The LED lamps also performed better in poor weather conditions compared to kerosene lamps. In addition to the low operating costs, fishers mentioned robustness of LED lamps as being among the reasons that inspired the shift. Some of the fishermen reported that on wavy nights, they have inadvertently submerged the LED lamps in water repeatedly without them being damaged. Thus, the robustness of the LED lamps allows them to perform better in wet and windy conditions compared to kerosene lamps, something that has not been noted specifically by other studies of artisanal fisheries (21-23).

Lamp type had only a minor effect on catch size. LED lamps resulted in slightly higher catches than kerosene lamps (Table 1), and units that were fishing by using LED-S and LED-R had slightly higher catches per unit effort compared to units equipped with kerosene lamps. The deeper depth of the LED light may be the reason for this slight impact on catch size. Catches did not differ between the two different LED lamp types, whether they were locally made from outdoor LEDs (LED-S) or a specialty LED for fishing (LED-R). Other studies examining a shift to LED light have mostly occurred in marine systems and typically find a large impact of using LED lamps relative to metal halide (24, 25) or compact fluorescent lamps (26). However, as was found here, the impact on efficiency appears to come primarily from fuel savings, whereas increases in net catch were typically minimal (24-26). An exception might be in a large Mediterranean lake, where the introduction of LED lamp rafts to replace kerosene lamps increased catch by 67% (22).

Lamp type did not have any effect on catch composition. The three species that were dominant in the fish catches are also dominant in the pelagic zone, where there are six species in total the four centropomids and two clupeids (4, 5). Even though their contribution was small, the other species that were caught are typically associated with near-shore or shallow environment (e.g. cichlids and sponges), consistent with the GPS coordinates provided by the fishermen showing fishing near the shoreline on some occasions (Fig 1). Sometimes the fishermen decide to fish nearshore because they believe the catch will be better and to save cruising fuel. In some cases, this decision is due to bad weather, and since our study period included months in the rainy season, strong storms could have occurred that affected their decision-making. Interestingly, the catches of small pelagic shrimp imply that nets were mended in such a way, potentially deliberately, that mesh size is reduced as the net is hauled up. Overall, lift net fishing closer to the shore results in greater catches of juveniles (3); however, the extent to which this could influence the overall fishery and lake ecology is complex (27).

Patchiness in pelagic fish catches is broadly recognized and is due to a range of factors, including environmental parameters, resource availability, predator-prey interaction, intraspecific relationships and habitat use (3, 28, 29). In our study, wind speed and moonlight were the main factors other than distance from the shore that affected fish catch. During high winds, rough water surface affects the beaming angle of light and refractions are more likely to happen. If the drifting speed is too high; fish may not stay with the lamp light. Moonlight counteracts the artificial lamp light and makes it less likely that fish congregate around artificial lamps placed close to the fishing nets, so fishermen generally do not fish during full moon (3, 4). The patchy spatial distribution of fishes (Fig 1) is reflected in the fact that the current variation in daily catches is only partially explained (21%) by the environmental variables we had included in our analysis.

Overall, we found that adoption of the new LED lamp technology on Lake Tanganyika was rapid and cost-effective, but the long-term environmental impacts were hard to discern. Our results concur with previous studies (23), indicating that increases in catch are small and the benefits of using LED lamps come primarily in terms of cost savings in kerosene fuel. If used in direct replacement of kerosene lamps, the observed transition to LEDs lamps may not have major effects on the fishery as long as the number of fishing boats remains constant. The LED lamps substantially reduce daily operating costs, but are not necessarily a clean technology, as city power or generators do most battery charging. Additionally, we recommend proper disposal of the lead-acid batteries being used by fishermen to power their lamps. Since fishermen will continue fishing during bad weather with the LED lamps (whereas with kerosene lamps they would not), future work could look at potential impacts on long-term fish catches and fishing pressure. Similarly, the cost efficiency of using LED lamps has potentially contributed to the emergent use of auxiliary boats with LED lamps, as well as an increase in the number of LEDs per unit, both of which could ultimately have an influence on catches.

## Acknowledgements

We thank Peter Limbu at The Nature Conservancy (TNC) for help with logistical arrangements and Omary Kashushu for help with field work and data collection, as well as staff at Tanzania Fisheries Research Institute for assistance and support. We thank the fishermen and boat owners for participating and the Beach Management Units at Kibirizi and Katonga and the municipal fisheries officers for allowing us to conduct the surveys and assisting with the selection of boat owners.

## Author contribution statement

HM, BMK, PBM, IK, CA, MN designed the study approach. HM, BMK, PBM, IK designed the experiment. BMK and HM wrote and administered the survey. HM, BMK, IK collected fisheries data, on which CL conducted QA/QC. PS and HM collected spectral data. BMK, HM, PS, IK conducted analyses. HM, BMK, and CMO made figures and tables. HM, BMK, and CMO wrote sections of the manuscript, and all authors contributed to revision and editing.

